# Synthetic Mucus Biomaterials for Antimicrobial Peptide Delivery

**DOI:** 10.1101/2023.03.07.531025

**Authors:** Sydney Yang, Gregg Duncan

## Abstract

Despite the promise of antimicrobial peptides (AMPs) as treatments for antibiotic-resistant infections, their therapeutic efficacy is limited due to the rapid degradation and low bioavailability of AMPs. To address this, we have developed and characterized a synthetic mucus (SM) biomaterial capable of delivering AMPs and enhancing their therapeutic effect. LL37 loaded SM hydrogels demonstrated controlled release of LL37 over 8 hours as a result of charge-mediated interactions between mucins and LL37 AMPs. Compared to treatment with LL37 alone where antimicrobial activity was reduced after 3 hours, LL37-SM hydrogels inhibited *Pseudomonas aeruginosa* PAO1 growth over 12 hours. LL37-SM hydrogel treatment reduced PAO1 viability over 6 hours whereas a rebound in bacterial growth was observed when treated with LL37 only. These data demonstrate LL37-SM hydrogels enhance antimicrobial activity by preserving LL37 AMP activity and bioavailability. Overall, this work establishes SM biomaterials as a platform for enhanced AMP delivery for antimicrobial applications.

## INTRODUCTION

Antibiotic resistance is a global health concern, resulting in over 5 million associated deaths worldwide in 2019 [1,2] The inability to completely eradicate bacteria at sites of infection has contributed to the rise in antibiotic–tolerant strains [3,4]. Within the United States, the Center for Disease Control estimates the economic burden of antibiotic resistant infections to be between $4-5 billion [5]. Recently, antimicrobial peptides (AMPs) have emerged as an attractive alternative to antibiotics. AMPs possess inherent antimicrobial activity and play important roles in host defense and innate immune response [6,7]. Cationic AMPs are capable of targeting the negatively charged bacteria cell membranes composed of mainly lipopolysaccharide [8]. Although, some bacteria and pathogens have evolved resistance mechanisms against AMPs by secreting proteases that target AMPs for proteolytic degradation [9,10]. In addition, AMPs are prone to degradation by host-derived extracellular proteases which result in rapid clearance times. As a result, the bioavailability and efficacy of AMPs is greatly reduced [11]. LL37, a human derived AMP approximately 37 residues in length, is cationic and α-helical in structure. These properties contribute to the broad spectrum antimicrobial activity of LL37 against numerous bacterial, viral, and fungal pathogens [12–14]. For *Pseudomonas aeruginosa*, the minimum inhibitory concentration (MIC) for LL37 is less than 10 μg/mL, indicative of high antimicrobial activity [12].

To address the limitations of AMPs, hydrogels and other biomaterial delivery systems have been utilized to protect AMP from degradation and extend their bioavailability [15,16]. Owing to the unique biological properties and barrier function of mucus, interest has grown in the design of mucin-based biomaterials for a diverse range of therapeutic applications [17,18]. Mucin glycoproteins, the main gel forming component in mucus, possess a wide profile of glycan domains which have been previously reported to modulate and dampen bacterial virulence [19,20]. In addition, negatively charged mucins have been reported to associate with cationic antimicrobial peptides [21]. This association of peptides to mucin is thought to protect the peptides from degradation [21,22]. Based on this evidence, we hypothesized mucin-based biomaterials and LL37 in combination would have a synergistic effect to provide long-lasting, enhanced antimicrobial activity.

Inspired by the native properties of mucus, we have previously designed a synthetic mucus (SM) hydrogel using commercially available porcine gastric mucins (PGM) and 4-arm polyethylene glycol (PEG) thiol [23]. However, a recent report suggests the processing of these mucins may lead to degradation of mucin glycans and other components that contribute to their native function [24]. Thus, we re-formulated SM hydrogels using mucins isolated in our lab using a gentler and scalable extraction procedure to preserve mucin-associated glycans and maximize its native activity. SM hydrogels were then utilized for encapsulating and delivering LL37 at concentrations ranging from near or above their previously reported MIC. We evaluated the physical, microrheological, release, and antimicrobial properties of the LL37 loaded SM hydrogels. The results of this study provide a basis to further investigate the utilization of mucus mimicking materials as tools for enhanced local AMP delivery and antimicrobial efficacy.

## MATERIALS AND METHODS

### Mucin extraction from native mucus

Porcine gastric mucins (PGM) used in formulating synthetic mucus (SM) hydrogels were extracted and purified using a method adapted from Sharma *et al* [25]. Fresh porcine small intestine was filled with 0.1M NaOH overnight to solubilize the mucus layer. The solution was then removed from the intestines and collected. To remove potential excess contaminants, the solution was reduced to a pH of 4 by adding 1M HCl. At a pH of 4, mucins will aggregate out of solution into a gel phase [25]. Afterwards, the solutions were centrifuged at 3500 rpm for 20 minutes to separate the aggregated mucins from solution. The supernatant was decanted, and the pellet was resuspended in deionized water. The pH of the solution was increased to 8 with 1M NaOH at which point the mucins will undergo a phase transition from gel to sol. The solution was stirred for 5 minutes, and the pH was altered to repeat the gel-sol cycle three times. DNase was added to the solution to a final concentration of 10 U/mL and incubated at 21°C overnight to remove excess DNA. The gel-sol cycle was repeated five times to further purify the mucin solution. After the last cycle, the solution was dialyzed in 100 kDa dialysis tubing (Spectra/Por) in deionized water. Dialyzed solutions were then transferred into conical tubes and frozen at -80°C overnight. Once frozen, the samples were lyophilized.

### Glycosylated mucin, protein, and DNA content

To determine glycoprotein (mucin) content, mucins were solubilized at 2% (w/v) in 50 μL of physiological buffer solution (154 mM NaCl, 3 mM CaCl_2_, and 15 mM NaH_2_PO_4_ at pH 7.4). A cyanoacetamide (CNA) reagent was made by mixing 200 μL of 2-cyanoacetamide with 1 mL of 0.15 NaOH. The mucin samples were mixed with 60 μL of CNA reagent and incubated at 100°C for 30 minutes. The fluorescence intensity of the samples was measured at an excitation/emission wavelength of 336/383 nm and compared to a standard curve produced from serially diluted solutions of bovine submaxillary mucin (BSM; Sigma-Aldrich) to determine the glycoprotein concentration. Protein concentration was quantified using a bicinchoninic acid (BCA) assay (Thermofisher) as described by the manufacturer. The glycoprotein to protein ratio was calculated to compare the mucin and protein content. To determine DNA content, 30 μL of 20% (w/v) 3,5-diaminobenzoic acid (DABA) solution was added to solubilized mucin samples and incubated at 60°C for 1 hour. To stop the reaction, 1 mL 1.76M HCl was added to the samples, and the fluorescence intensity was measured at an excitation/emission wavelength of 390/530 nm.

### Preparation of LL37-SM hydrogels

4% (w/v) PGM were mixed in a physiological buffer solution (154 mM NaCl, 3 mM CaCl_2_, and 15 mM NaH_2_PO_4_ at pH 7.4). Salts within the buffer help to solubilize PGM in solution [23]. LL37 (Anaspec) was directly added into the PGM solution and stirred for 2 hours at room temperature to ensure thorough mixing. 8% (w/v) 4-arm PEG-thiol was prepared in physiological buffer in a separate vial. The PGM-LL37 and PEG-thiol solutions were mixed at a 1:1 ratio to synthesize the AMP-SM hydrogel solution at a final composition of 2% (w/v) PGM and 4% (w/v) PEG-thiol. To make hydrogel discs, 100 μL of the mixed LL37-SM solution was transferred to cylindrical molds. The LL37-SM hydrogel solution was incubated for 12 hours at room temperature to allow for gelation. LL37-SM hydrogels were prepared with 100, 50, and 10 μg/mL LL37 as final concentrations.

### Hydrogel degradation and swelling

To evaluate degradation and swelling over time, LL37-SM hydrogels were prepared as 100 μL discs and submerged in 1 mL of phosphate buffer saline (PBS, Sigma-Aldrich) and incubated at 37°C. At hour timepoints over 24 hours, the PBS was discarded and the masses of the hydrogels were measured to determine extent of degradation. After 4 hours of incubation, the swelled mass, (*M*_s_) of the hydrogels was measured. The hydrogels were then lyophilized and the dried mass (*M*_d_) was measured. Swelling ratio, defined as the fractional increase in mass due to water absorption, was calculated using **Eq. 1**.

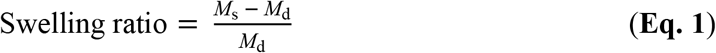

### Preparation of nanoparticles for multiple particle tracking microrheology

Carboxylate-modified fluorescent polystyrene nanoparticles (NP; Life Technologies) of 100 nm diameters were coated with polyethylene glycol (PEG) to form high-surface-densities. PEG conjugation was completed via carboxyl-amine linkage using 5 kDa methoxy PEG-amine (Creative PEGWorks) as previously reported [23,26]. Zeta potential of the NPs was measured in 10 mM NaCl at pH 7 using a NanoBrook Omni (Brookhaven Instruments).

### Multiple particle tracking microrheological characterization of LL37-SM hydrogels

The rheological properties of the LL37-SM hydrogels were characterized using particle tracking microrheology (PTM) [27]. Microscopy chambers were made of o-rings coated with vacuum grease on slides. To prepare samples for the microscopy chamber, 25 μL of LL37-SM hydrogel was added to the microscopy chamber and 1 μL of ∼0.002% w/v suspension PEG-coated NPs was mixed into the hydrogel sample. The hydrogels were allowed to equilibrate at 21°C for 30 minutes. NP diffusion through the hydrogels was evaluated via fluorescence video microscopy as described in our work previously [23]. NP trajectories were measured via a series of videos taken using fluorescent microscopy (Zeiss Confocal LSM 800) fitted with a 63x water-immersion objective and analyzed using a previously developed algorithm [23,28]. Using the trajectories, the ensemble average mean squared displacement (MSD) as a function of lag time (τ) was calculated as MSD= ([*x*(*t*+τ)-*x*(*t*)]^2^+[*y*(*t*+τ)-*y*(*t*)]^2^). The logarithm (base 10) of MSD at a time scale of τ = 1s (log_10_[MSD_1s_]) was used to measure diffusion rate. The loss modulus (*G”)* was calculated from the Laplace transformation of (MSD(τ), MSD(s)) using the generalized Stokes-Einstein equation *G*” = 2*k*_B_*T*/[π*as*MSD], where *k*_b_*T* is the thermal energy, *a* is the particle radius, and *s* is the complex Laplace frequency [29]. The complex modulus *G*\* was obtained as *G*\*(ω) = *G*’(ω) + *G*’’(iω), where iω is substituted for *s, i* is a complex number, and ω is frequency. From the determined *G\**, the complex microviscosity (η*) was calculated using η* = *G*\*(ω)/ ω and evaluated at a frequency of ω = 1 Hz [23].

### Release of LL37 from SM hydrogels

LL37 loaded SM hydrogels were made in cylindrical molds to create 100 μL hydrogel discs with varying concentrations of fluorescently labelled tetramethylrhodamine-LL37 (TRITC-LL37, Rockland). A standard curve of TRITC fluorescence intensity as a function of LL37 concentration was generated to enable the quantification LL37 release. The LL37-SM hydrogel discs were submerged in 1 mL of PBS and incubated at room temperature of 37°C. At timepoints of 2, 4, 6, 12, and 24 hours, the PBS was collected and replaced with 1 mL of fresh PBS. To determine the concentration of LL37 released, TRITC fluorescence within the collected PBS samples were quantified at an excitation wavelength of 557 nm and emission wavelength of 576 nm. LL37 concentration was calculated from the TRITC fluorescence intensity and used to evaluate release profiles plotted as fluorescence intensity over time. From this curve, the release exponent *n* was calculated via the Korsmeyer-Peppas model (**Eq. 2**), which is used to describe drug release from cylindrical polymeric systems [30]. The Korsmeyer-Peppas model is defined as:

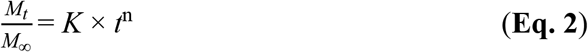

where *M*_t_ is the mass released at time *t, M*_∞_ is the mass released at time ∞, *K* is the release rate constant, *t* is time, and *n* is the release exponent. This process was repeated for LL37 loaded SM hydrogels in 200 ng/mL trypsin, representative of trypsin concentration in serum [31].

### Release kinetics of LL37 in SM hydrogels

Fluorescence recovery after photobleaching (FRAP) was used to assess the release kinetics and diffusion characteristics of LL37 through the SM hydrogel. LL37-SM hydrogel solutions were made with concentrations of TRITC LL37. Microscopy chambers were prepared with 25 μL of hydrogel. To conduct FRAP, we used fluorescence confocal microscopy (Zeiss Confocal LSM 800) with a 63x water-immersion objective. Prior to bleaching, 10 images were taken. LL37-SM hydrogels were bleached with a 561 nm laser using a circular region of interest (ROI) with a diameter of 20 μm at 100% laser transmission at 40 iterations. After bleaching, images were captured every 10 milliseconds over 100 seconds using a 561 nm laser at 10% intensity.

### Planktonic PAO1 growth with LL37-SM hydrogel treatment

LL37-SM hydrogel discs were made with concentrations of LL37. Planktonic green fluorescent protein (GFP) expressing PAO1 (ATCC 10145GFP) were cultured in MOPS minimal media (Teknova). An optical density of 0.1 at 600 nm (OD_600_) was used as the working concentration of bacteria. After measuring the optical density, the LL37-SM hydrogel disc was then placed into 1 mL of PAO1 culture and incubated at 37°C. At designated hour timepoints from 1-24 hours, 100 μL of media was collected and replaced with fresh media. GFP fluorescence intensity of the collected media was measured at 510 nm.

### Planktonic PAO1 viability with LL37-SM hydrogel treatment

Planktonic PAO1 (ATCC 15692) was cultured in lysogeny broth (LB, Sigma-Aldrich) to a working concentration at an optical density of 0.1 at OD_600_. As described earlier, 100 μL LL37-SM hydrogel discs were prepared and placed in 1 mL of bacteria culture and incubated for 3 hours at 37°C. After incubation, the bacteria culture media was collected and made into serial dilutions of 1×10^5^ to 1×10^8^ for spot plating 10 μL spots onto agar plates. The colony forming units (CFUs) were counted after the plates were incubated at 37°C for 24 hours. Based on the plated volume, the logarithm (base 10) of CFU/mL was calculated and used to quantify PAO1 viability.

### STATISTICAL ANALYSIS

Data were graphed and statistically analyzed using GraphPad Prism 9. Bar graphs show means with standard deviations, box and whisker plots show medians with 5th to 95th percentiles, and line graphs show means with standard deviations. Unpaired t-test was performed for analysis between two groups. For analysis between multiple groups, a one-way ANOVA with Tukey’s post hoc correction was performed. A two-way ANOVA was conducted to compare multiple groups with two independent factors. Kruskal-Wallis test with Dunn’s comparison was used to compare multiple groups with non-Gaussian distributions. Values of *P*<0.05 were considered statistically significant.

## RESULTS AND DISCUSSION

### Extraction of mucins from native mucus

We have adopted a method inspired by Sharma *et al* to extract porcine gastric mucins (PGM) from native porcine small intestine mucus [25]. The mucus layer is completely solubilized, and through alternating the pH, the aggregation of mucins in solution can be controlled. At a pH of 7 and above, PGM is electronegative and hydrophobic domains remain enfolded within the mucin structure. At a pH of 4, mucins unfold and expose previously buried hydrophobic domains. Hydrophobic interactions between the hydrophobic domains promote mucin cross-linking, allowing mucins to undergo a sol-gel transition [32]. Centrifugation of the gel phase enables the collection of aggregated mucins and removal of cellular contaminants (**Figure 1A**). The extracted PGM contained approximately 30% residual DNA, a higher quantity than that found in commercial PGM (**Figure 1B**). However, compared to prior literature, similar mucin extraction methods have resulted in comparable DNA content [25]. The glycosylated mucin and protein contents were measured and used to determine the mucin to protein ratio by mass. We determined a significant increase in the mass protein content in extracted mucins (**Figure 1C)**. There was no significant difference between the mucin to protein ratios between commercial PGM and our extracted PGM (**Figure 1D**). Excess residual DNA may be the result of DNA binding to positively charged amino acids on non-glycosylated domains that become exposed during the sol phase and remain entangled within the mesh of mucin molecules during the gel phase [33]. Commercial preparation of mucins includes the treatment of mucus with pepsin and additional proteases to isolate mucins. However as noted, protease treatment has been shown to fragment mucins and alter mucin functionality [24]. Overall, this procedure provides a highly scalable method to extract mucins that are minimally altered and were used in this work to formulate synthetic mucus hydrogels.

**Figure 1.**
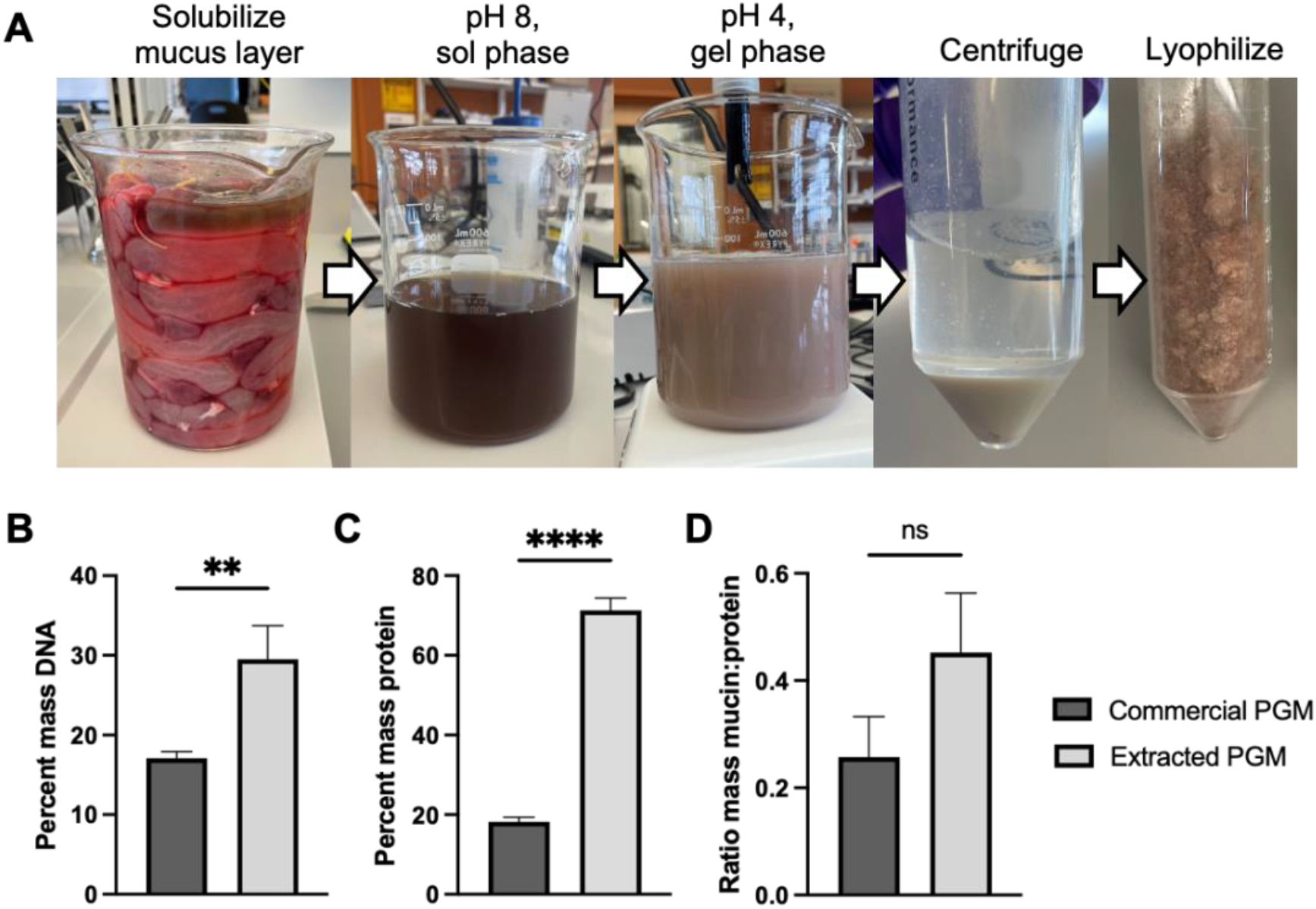
Extraction of porcine gastric mucins (PGM). (**A**) Schematic of preparation of mucin extraction and preparation from porcine small intestine tissue. (**B**) DNA content present in commercially available and in-house extracted PGM (*n* = 3). (**C**) Glycosylated mucin to protein ratio of content in commercial and extracted PGM (*n* = 3). \*\**P*<0.01 and \*\*\*\**P*<0.0001 for unpaired t-test.

### Engineering synthetic mucus hydrogels for LL37 encapsulation

We have previously reported the design of a synthetic mucus hydrogel using 4-arm PEG thiol to form disulfide bonds and enable hydrogel formation [23]. Based on this design, we prepared synthetic mucus (SM) hydrogels with 2% (w/v) in-house extracted PGM and 4% (w/v) PEG-4SH to mediate crosslinking (**Figure 2A**). Mucins are negatively charged macromolecules and have been previously shown to bind to cationic peptides via non-specific electrostatic interactions [21,22]. Based on this charge mediated interactions between mucins and LL37 AMPs, we hypothesized LL37 will associate to mucin glycoproteins and slowly release over time from our hydrogel formulation. When LL37 is encapsulated in the SM hydrogel, we will refer to them as LL37-SM hydrogels throughout the article.

**Figure 2.**
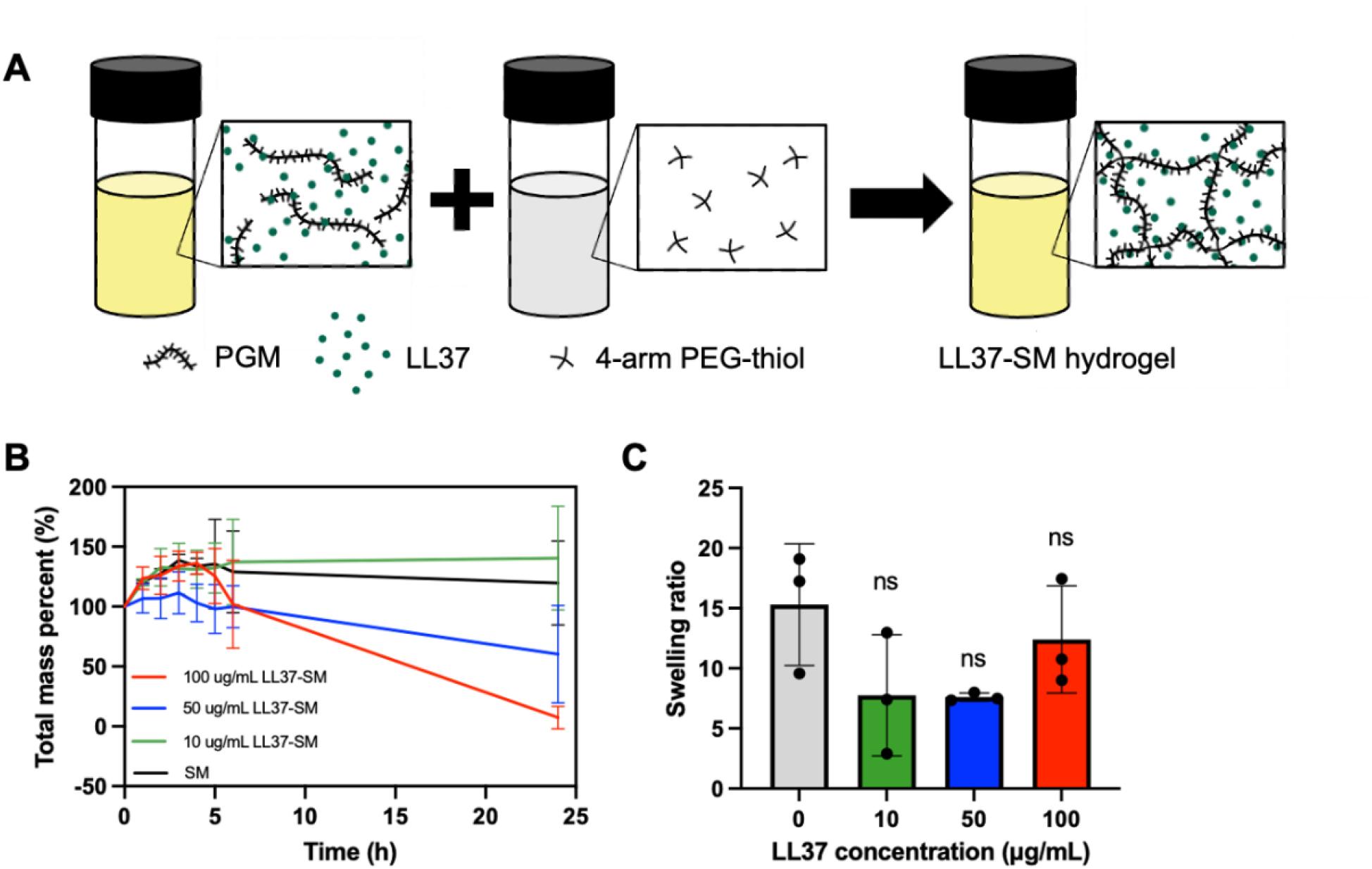
Physical characterization of LL37-SM hydrogels. (**A**) Schematic of LL37-SM hydrogel preparation. (**B**) Degradation profile of LL37-SM hydrogels over 24 hours (*n* = 3). (**C**) Swelling ratio of LL37-SM hydrogels after 4 hours in PBS (*n* = 3). No statistical significance observed by one-way ANOVA.

To characterize the degradation of the LL37-SM hydrogels, we formulated the hydrogels as discs and submerged the discs into PBS to observe the change in mass over time. The degradation profile showed all SM hydrogels initially swelling across 4 hours. Beyond 4 hours in PBS, the SM and 10 μg/mL LL37-SM hydrogels portrayed little to no decrease in mass over 24 hours. Meanwhile, the 50 and 100 μg/mL LL37-SM hydrogels exhibited a more rapid decrease in mass, and therefore, a greater rate of degradation over 24 hours (**Figure 2B**). Thus, it appears increasing the LL37 concentrations within SM hydrogels increased the rate of degradation. Increasing concentration of LL37 within the hydrogel may affect the net osmotic pressure which may contribute to the faster degradation observed. However, the mechanism by which this occurs has not been determined. Nonetheless, the gradual hydrogel degradation observed over 24 hours is expected to aid in the slow and prolonged release of LL37 over time. In addition, the degradation profiles demonstrate SM hydrogels, when loaded with higher concentrations of LL37, degrade completely in PBS, which may potentially aid in the complete release of LL37. The swelling ratios were determined after 4 hours of swelling in PBS and were not significantly different among all SM hydrogels (**Figure 2C**). All SM hydrogels swell upwards to an estimated 7 to 15 times the hydrogel dry mass. Hence, SM hydrogels have a high water affinity and would potentially improve water retention as a topical application at local sites.

### LL37-SM hydrogels form elastic dominant gels

Using PTM, we assessed the rheological properties at the micro- and nano-scale to locally resolve the rheological moduli and microviscosity [27]. As compared to traditional rheological characterization, we were also able to examine biophysical properties using only small quantities of these biomaterials which enabled us to evaluate a range of LL37 loading concentrations more easily. Increasing LL37 concentration within SM hydrogels showed a statistically significant decrease in nanoparticle diffusion as indicated by measured log_10_[MSD_1s_] (**Figure 3A**). Additionally, the 50 and 100 μg/mL LL37-SM hydrogels showed a significant decrease in log_10_[MSD_1s_] compared to the 10 μg/mL LL37-SM hydrogel. The MSD data indicates nanoparticle diffusion within the hydrogel mesh network has decreased when the LL37 concentration is increased. An increase in cross-linking density and mesh entanglement promoted by the electrostatic interactions between LL37 and the mucin network may contribute to the decrease in MSD observed. All hydrogels showed *G’/G”* ratios greater than 1, indicating the hydrogels were elastic dominant. No significant difference was observed among the *G’/G”* ratios for different LL37 concentrations within SM hydrogels (**Figure 3B**). Thus, the LL37-SM hydrogels have comparable moduli ratios to that of SM hydrogels. In addition, the 50 and 100 μg/mL LL37-SM hydrogels demonstrated a significant increase in complex microviscosity (*η*)* compared to the SM and 10 μg/mL LL37-SM hydrogels (**Figure 3C**). The PTM data confirmed our SM formulation forms hydrogels when loaded with LL37.

**Figure 3.**
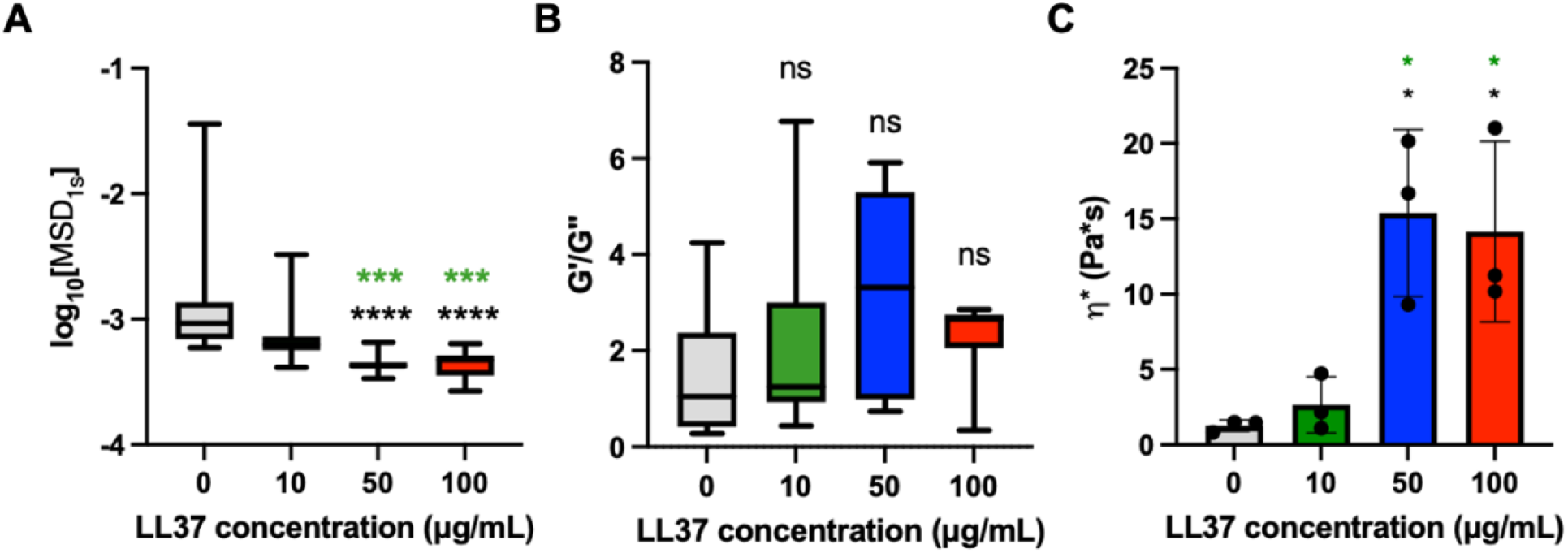
Microrheological characterization of LL37-SM hydrogels. (**A**) Measured log_10_(MSD_1s_) of 100 nm PEG-coated NP in LL37-SM hydrogels after 24 hours of incubation at room temperature (*n* = 20). \*\*\**P*<0.001 and \*\*\*\**P*<0.0001 for Kruskal-Wallis test. (**B**) Elastic and viscous moduli ratio (*G’/G”*) at a frequency ω = 1 Hz calculated from measured MSD (*n* = 6). No statistical significance observed by Kruskal-Wallis test. (**C**) Mean complex microviscosity (*η\**) at a frequency ω = 1 Hz calculated from measured MSD (*n* = 3). \**P*<0.05 for one-way ANOVA. The color of the asterisk indicates the comparison group.

### SM hydrogels show gradual release of and high association with LL37

Fluorescence recovery after photobleaching (FRAP) was used to evaluate the diffusion and association of LL37 within SM hydrogels. From the FRAP fluorescence recovery profiles, the mobile fraction, based on relative fluorescence intensity at t = 100 s, were determined to be 0.20, 0.30, and 0.35 for 10, 50, and 100 μg/mL LL37-SM hydrogels, respectively (**Figure 4A**). These data indicate LL37 is highly associated with the SM hydrogel as we expected with a majority of LL37 immobilized in the SM hydrogel. This high association of LL37 is hypothesized to be driven by binding via electrostatic interactions between positively charged LL37 and negatively charged mucin molecules. At lower concentrations of LL37, there may be more binding sites available on mucins for LL37 to interact with. However, as the LL37 concentration increases, the available binding sites may become limited and eventually reach a saturation threshold in which less LL37 is bound to mucin.

**Figure 4.**
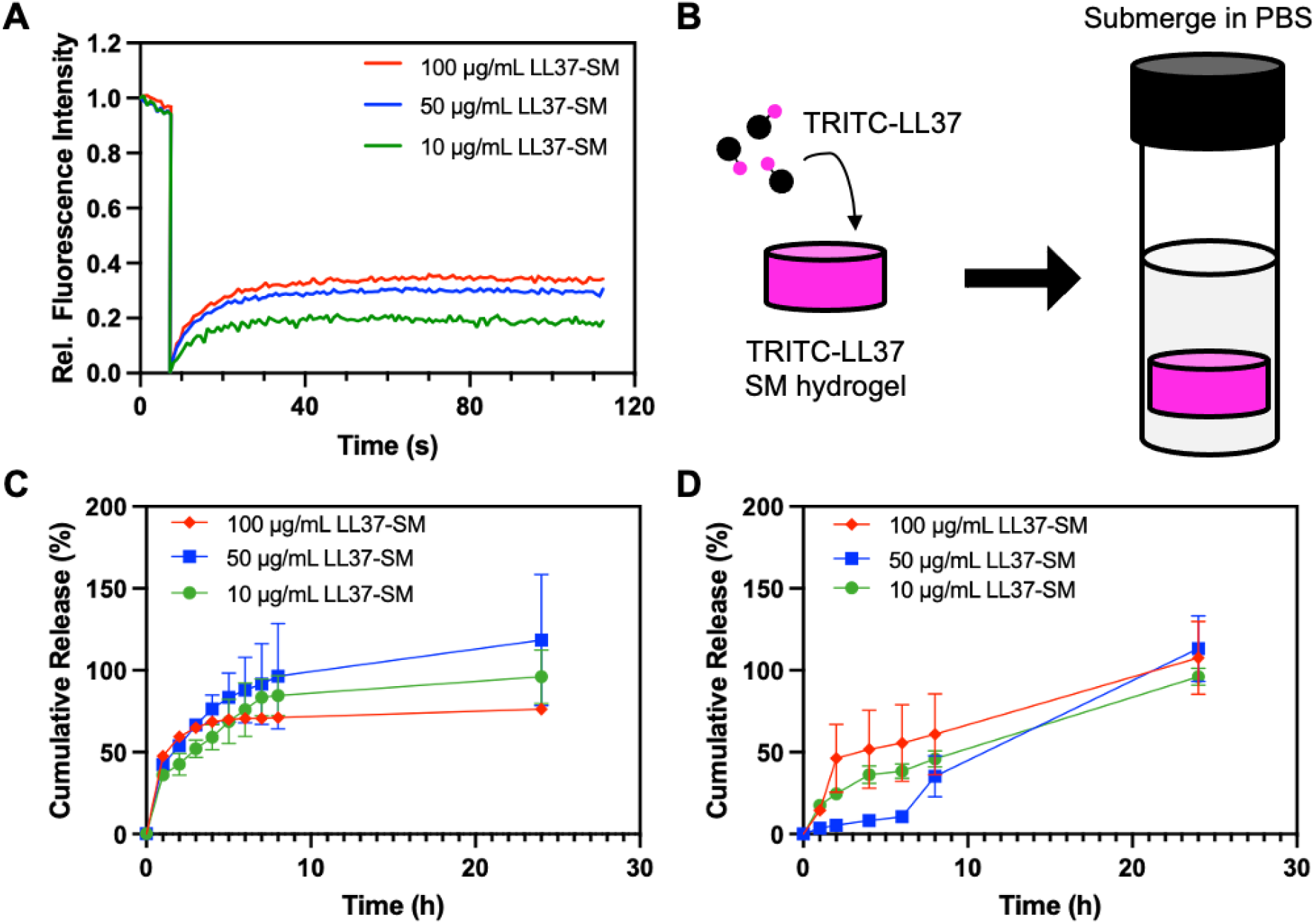
Release kinetics of LL37 from SM hydrogels. (**A**) FRAP mean relative fluorescence intensity profile of LL37-SM hydrogels (*n* = 3). (**B**) Schematic of TRITC-LL37-SM hydrogel set up for evaluating release. Cumulative mass release profiles of LL37 from SM hydrogels (*n* = 3) (**C**) without and (**D**) with 200 ng/mL trypsin over 24 hours (*n* = 3).

We characterized the release properties of the LL37-SM hydrogels by measuring the release of TRITC LL37 from SM hydrogel discs submerged in PBS (**Figure 4B**). SM hydrogels demonstrated initial burst release over 6 hours and maximum release LL37 over 8 hours. Approximately 90%, 95%, and 70% cumulative mass release was measured for 10, 50, and 100 μg/mL LL37-SM, respectively (**Figure 4C**). The remaining LL37 not released may be bound to mucin or unable to diffuse through the hydrogel mesh network. From the LL37 concentrations assessed, the upper limit for mass release from our SM hydrogel discs was approximated to 7 μg LL37.

In addition, the release profiles were measured with the treatment of trypsin, a protease expressed at infection and inflammation sites and known to degrade mucins, to assess the effect on LL37 release [34–36]. LL37-SM hydrogels demonstrated a slower release of LL37 when treated with trypsin (**Figure 4D**). However, LL37 release does not appear to be dependent on LL37 concentration. Trypsin is expected to rapidly SM hydrogel as trypsin able to cleave the mucin peptide backbone. Previously reported, colonic mucins treated with trypsin resulted in fragments comprised of decreased concentrations of positively charged amino acids and increased concentrations of hydrophobic residues [35]. The reduction of positively charged residues, may decrease repulsion between mucin domains and promote greater LL37 to mucin binding. In addition, the cleavage of mucin may expose more available sites for LL37 to bind through both electrostatic and hydrophobic interactions, and therefore, result in slower release. However, we must also consider the degradation of LL37 by trypsin [37–39]. Additional characterization of release kinetics for other AMP and peptide cargo from SM in future work will help to elucidate what drives the observed slowing of release in response to proteolysis.

### Fitting of LL37 release to the Korsmeyer-Peppas model

To compare and assess the release mechanism of LL37-SM hydrogels, we fitted the release profiles to the Korsmeyer-Peppas model which describes drug release from cylindrically–shaped polymeric systems [30]. Using the Korsmeyer-Peppas model, we determined the values for the release exponent, *n*, which is used to describe the release mechanism. For a cylindrical geometry, values of *n*≤0.45 indicate a diffusion dominant release, 0.45<*n*<1 indicate anomalous, non-Fickian diffusion transport, and *n*≤1 indicate anomalous, time independent release [30]. Release of LL37 from the SM hydrogel was determined to be Fickian diffusion dominant (**Table 1**). In the presence of trypsin, release is anomalous and non-Fickian diffusion dominant as expected, supports the observed release profiles, and suggests hydrogel degradation plays an important role in LL37 release in the presence of trypsin. Additionally, the time required for 50% cumulative release of LL37, or the half-time *t*_*1/2*_, was calculated and ranged from approximately 0.5 to 2.6 hours and 5 to 21 hours without and with trypsin treatment, respectively. The half-times for LL37 release did not appear to be concentration dependent. Increased half-time and prolonged release is expected to increase LL37 bioavailability. Interestingly, the half-times for LL37-SM release with trypsin are greater than the half-life of LL37 in PBS, which is estimated to be 0.5 - 1 hours [40]. Therefore, treatment with trypsin decreases the rate of LL37 release, and together with the high association of LL37 to SM, benefit in prolonging LL37 half-life and bioavailability.

**Table 1.** Release profiles were fitted to the Korsmeyer-Peppas model. *R*^*2*^ values, release exponents (*n*), release mechanisms, and half-times (*t*_*1/2*_) were determined for LL37-SM hydrogels with and without the addition of 200 ng/mL trypsin.

| Condition, LL37-SM | $R^2$ | Release exponent, $n$ | Release mechanism | Half-time, $t_{1/2}$ (h) |
| --- | --- | --- | --- | --- |
| 10 $\mu\text{g/mL}$ | 0.78 | $0.39 \pm 0.03$ | Diffusion | 2.60 |
| 10 $\mu\text{g/mL}$ + trypsin | 0.91 | $0.46 \pm 0.09$ | Anomalous | 9.61 |
| 50 $\mu\text{g/mL}$ | 0.89 | $0.42 \pm 0.08$ | Diffusion | 0.51 |
| 50 $\mu\text{g/mL}$ + trypsin | 0.76 | $0.93 \pm 0.31$ | Anomalous | 21.0 |
| 100 $\mu\text{g/mL}$ | 0.58 | $0.24 \pm 0.13$ | Diffusion | 1.11 |
| 100 $\mu\text{g/mL}$ + trypsin | 0.99 | $0.60 \pm 0.31$ | Anomalous | 4.96 |

### LL37-SM hydrogels inhibit Pseudomonas aeruginosa growth

We evaluated the antimicrobial activity of LL37-SM hydrogels on planktonic *P. aeruginosa* PAO1 (**Figure 5A**). To determine the effect on growth, we treated GFP expressing PAO1 cultures with 100 μg/mL LL37-SM hydrogels. Compared to the untreated PAO1, treatment with free LL37 inhibited growth over 12 hours (**Figure 5B**). Meanwhile, treatment with SM and LL37-SM hydrogels exhibited bactericidal activity over the first 2 hours and continued to inhibit growth over 12 hours. Interestingly, the SM hydrogels alone without AMP demonstrated a similar trend in reducing PAO1 growth. It should be noted PGM used in our SM formulation is a crude extract and does contain excess proteins. Thus, the PGM extract consists of various proteins and peptides which may possess antimicrobial activity as they are naturally present within the gastrointestinal tract [41,42]. This likely explains the intrinsic antimicrobial activity observed for SM hydrogels.

**Figure 5.**
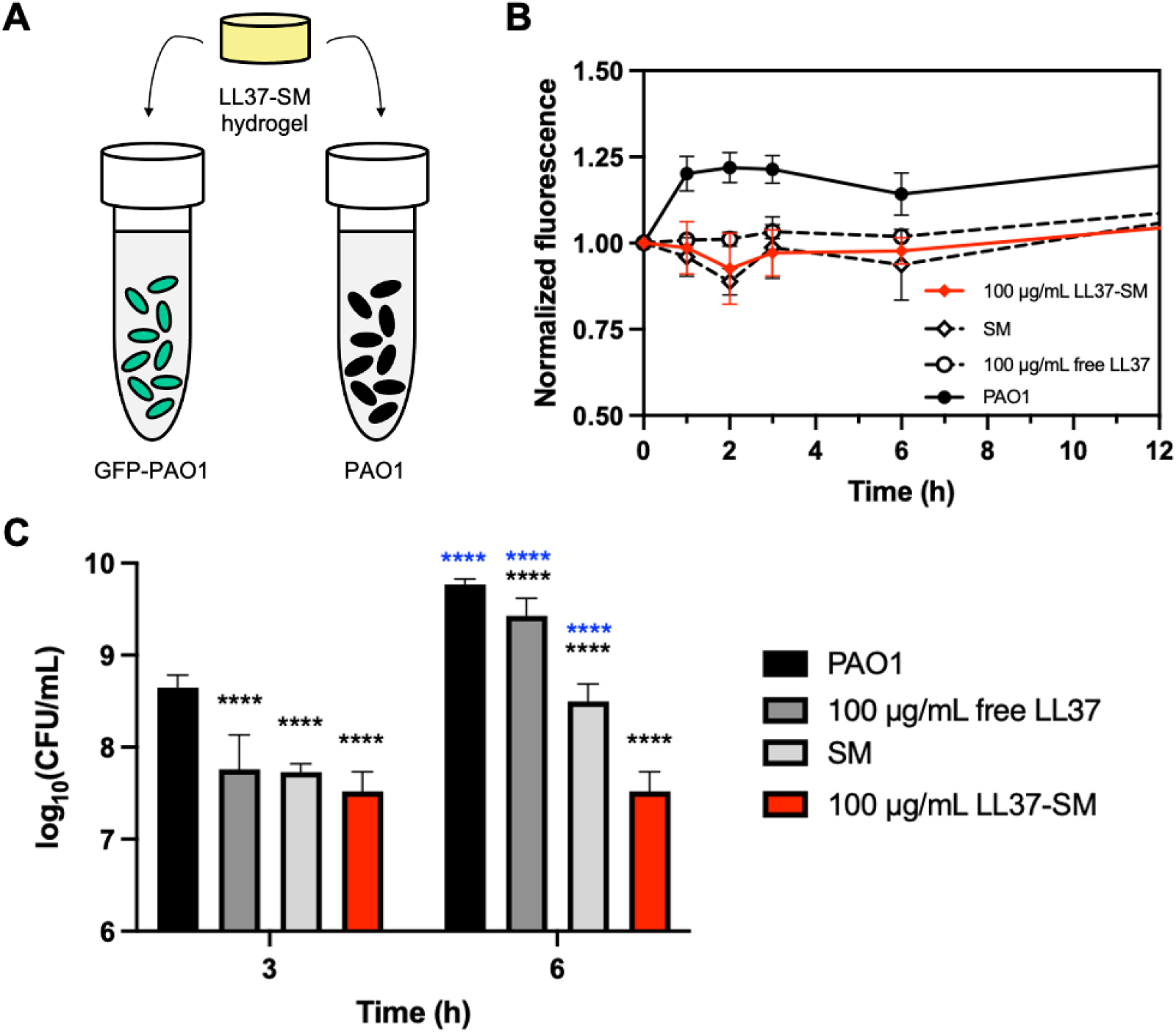
Antimicrobial activity of LL37-SM hydrogels against planktonic PAO1. (**A**) Schematic of hydrogel treatment on planktonic GFP-PAO1 and PAO1 to assess the effect on growth and viability, respectively. (**B**) Growth profiles of planktonic PAO1 with treatment (*n* = 3). (**C**) PAO1 viability after 3 hours of treatment (*n* = 3). \*\*\*\**P*<0.0001 for two-way ANOVA. Blue asterisks indicate the comparison between time groups.

The viability of PAO1 after treatment was assessed via CFU count by plating. Treatment with free LL37, SM, and LL37-SM hydrogels significantly reduced PAO1 viability after 3 hours of treatment (**Figure 5C**). After 6 hours of treatment, free LL37 and both SM and LL37-SM hydrogels showed significantly reduced PAO1 viability compared to the untreated control. However, the free LL37 and SM hydrogel treatment showed a significant increase PAO1 viability from 3 to 6 hours. Only the LL37-SM hydrogel demonstrated continued inhibition of PAO1 viability across 6 hours of treatment. This suggests that without LL37, SM hydrogels have a very short-lived antimicrobial effect, and with LL37, SM hydrogels exhibit a synergistic effect. In addition, the SM hydrogels preserve the antimicrobial activity of LL37 across 6 hours, which also suggests prolonged bioavailability for treatment.

## CONCLUSION

We have developed a mucus-inspired biomaterial for the encapsulation and delivery of AMPs. Treatment with trypsin resulted in overall slower LL37 AMP release which may provide future interest to design protease responsive degradation for controlled release. Encapsulation of LL37 by SM is expected to be driven by electrostatic interactions, in which LL37 demonstrated high association to SM. As such, SM provides a modular platform for delivery of cationic AMPs enabling its use for a wide number of infectious disease applications. LL37-SM hydrogels have enhanced antimicrobial therapeutic action with sustained inhibition of bacteria growth and viability. In conclusion, this work establishes a promising new biomaterial strategy which will be tested further in pre-clinical *in vivo* infection models.

## ACKNOWLEDGEMENTS

This study was supported by the NIH (EB030834 awarded to G.A.D.), the NSF GRFP (DGE1840340 awarded to S.Y.), and the University of Maryland. S.Y. is supported by the NSF GRFP and the University of Maryland Clark Fellowship.

